# Overcoming air-water interface-induced artifacts in Cryo-EM with protein nanocrates

**DOI:** 10.1101/2025.08.18.667046

**Authors:** Matthew C Jenkins, Daija Bobe, Jake D Johnston, Jonah Cheung, Akira Karasawa, Christina M Zimanyi, Ömer Dermanci, M.G. Finn, Alex de Marco, Mykhailo Kopylov

**Affiliations:** School of Chemistry and Biochemistry, School of Biological Sciences, Georgia Institute of Technology, Atlanta, GA, 30332, USA; Simons Electron Microscopy Center, New York Structural Biology Center, New York, NY, 10027, USA; Department of Physiology and Cellular Biophysics, Columbia University, New York, NY, 10027, USA

## Abstract

Contact with the air-water interface can bias the orientation of macromolecules during cryo-EM sample preparation, leading to uneven sample distribution, preferred orientation, and damage to the molecules of interest. To prevent this, we describe a method to encapsulate target proteins within highly hydrophilic, structurally homogeneous, and stable protein shells, which we refer to as “nanocrates” for this purpose. Here, we describe packaging, data acquisition, and reconstruction of three proof-of-principle examples, each illuminating a different aspect of the method: apoferritin (ApoF, demonstrating high-resolution), thyroglobulin (Tg, solving a known preferred orientation problem), and 7,8-dihydroneopterin aldolase (DHNA, a structure previously uncharacterized by cryo-EM).

## Main

Cryo-electron microscopy (cryo-EM) can provide high-resolution structural information about biomolecules in vitreous ice. Resolving such three-dimensional reconstructions requires averaging of tens to hundreds of thousands of randomly oriented individual particles. However, biomolecules frequently concentrate at the air-water interface (AWI) during sample preparation, often adopting preferred orientations that make three-dimensional reconstruction difficult or impossible due to insufficient orientation sampling.^1,2^ Several methods have been developed to address this issue,^3-5^ including sample tilting during data collection,^6^ modifications to EM grids^7-9^ (often involving the addition of randomly-oriented binding interactions^10-12^), changes in sample preparation or freezing methods,^7,13-16^ and the use of detergents^17-19^ or barrier proteins.^20,21^ DNA superstructures have also been used to orient encapsulated or attached proteins for cryo-EM analysis, although for different purposes, resulting in reconstructions of moderate resolution.^22,23^ None of these methods offers a complete solution to AWI-related issues.

Our approach was inspired by the established use of natural virus capsids and engineered protein containers for packaging applications.^24-27^ We adapted this concept for cryo-EM by encapsulating target proteins within symmetric MS2 bacteriophage-derived “nanocrates”. Being highly symmetric, the nanocrates do not adopt a preferred orientation at the AWI, and therefore their enveloped cargoes are randomly oriented. Nanocrate densities can be computationally subtracted during single-particle cryo-EM analysis to allow for high-resolution reconstruction of the molecules inside.

MS2 virus-like particles have multiple properties that make them ideal for use as nanocrates. They are expressed and isolated in high yields and are amenable to controlled disassembly and reassembly around cargo proteins by a simple pH-dependent protocol.^28-30^ Disassembled MS2 coat proteins can be stored at 4^°^C for at least a day and at -80^°^C for several months without losing reassembly and encapsulation efficiency. At 21 nm, the interior diameter of the MS2 particle is large enough to accommodate many target cargoes.^31^ Importantly, the MS2 nanocrates themselves are highly monodisperse (97% of wild-type MS2 particles are T=3 icosahedra),^31,32^ which is a requirement for the signal subtraction step needed to reconstruct cargoes.^33-35^ As expected, packaged cargo molecules are randomly oriented with respect to the cryo-EM grid, enabling high-resolution isotropic reconstructions, as demonstrated here for three proteins at resolutions of 2.1-2.9 Å.

We first packaged apoferritin (ApoF)—a homo 24-mer of 484 kDa total molecular weight, often used as a standard cryo-EM high-resolution benchmark (**Figure 1A**). Briefly, we purified MS2 capsids from *E. coli*, performed disassembly in an acidic solvent with the removal of packaged viral RNA by centrifugation, and reassembled the capsid proteins at neutral pH in the presence of ApoF. We designate the reassembled cages as nanocrates (“nc”) to distinguish particles derived from disassembly and reassembly, rather than another commonly used packing strategy in which cargo-containing virus-like particles (VLPs) are derived from simultaneous expression and packaging in the expression host.^24,36,37^ The @ symbol is used to designate the packaged species. After reassembly, the reaction was concentrated 10-fold, vitrified on cryo-EM grids, and imaged on a Thermo Fisher Scientific Titan Krios microscope equipped with a Gatan K3 camera.

**Figure 1.**
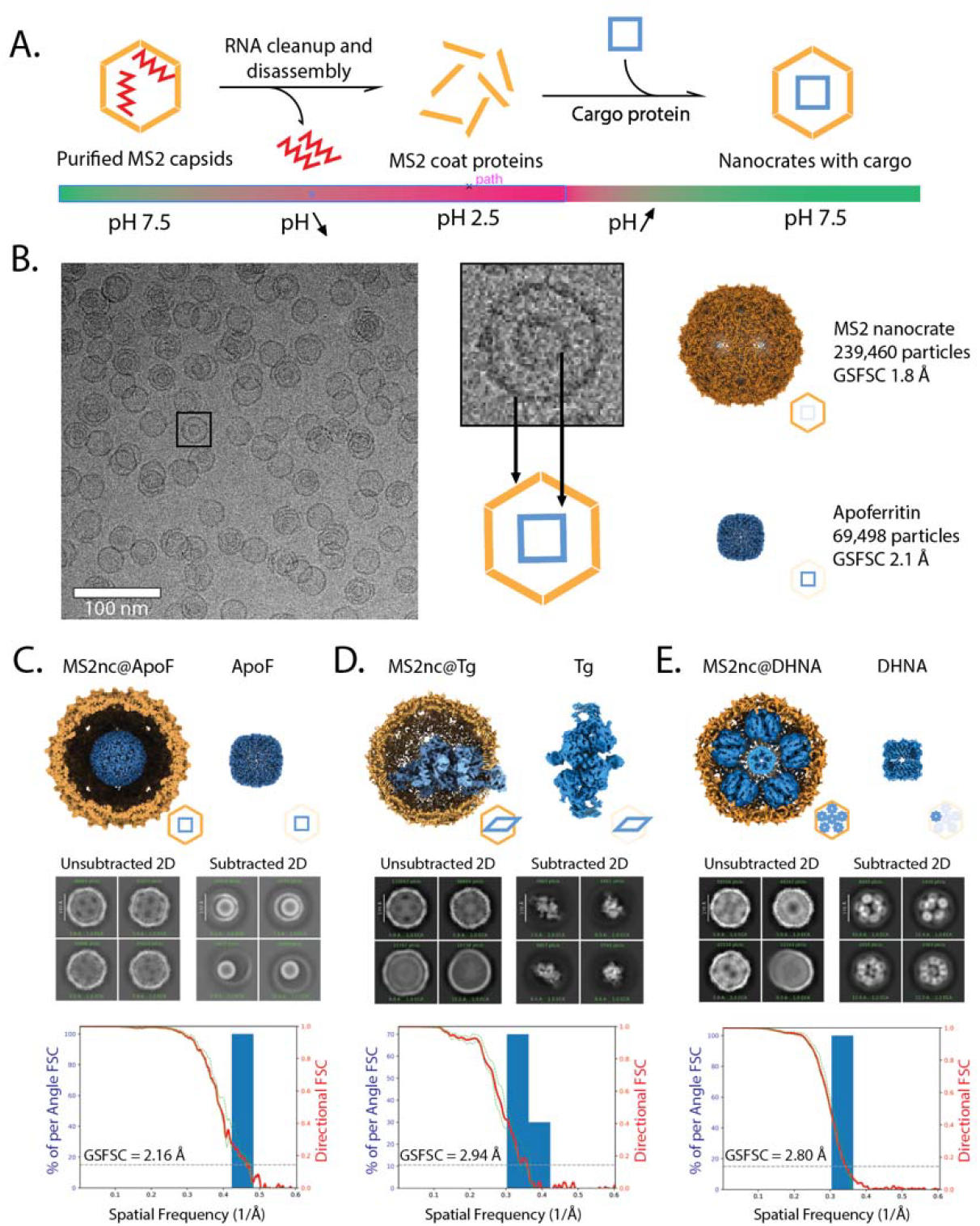
Structure determination of three proteins from nanocrates. **A**. Schematic representation of the nanocrate method as described in the text. **B**. Representative cryo-EM micrograph of MS2nc@ApoF particles, with a cropped out single particle showing the MS2 shell and encapsulated ApoF cargo and cryo-EM maps obtained after processing the MS2nc@ApoF dataset. The MS2nc was processed with icosahedral symmetry; after particle subtraction, ApoF was processed with octahedral symmetry. **C-E**. Summary of nanocrate-based structure determination for ApoF, Tg, and DHNA, respectively. (*Top*) Complete reconstructions, with cargo proteins in blue and MS2nc in yellow shown to the left; cargo-only reconstructions are shown on the right. (*Middle*) 2D class averages before and after nanocrate density subtraction. (*Bottom*) 3D FSC plots for each cargo protein.

Cryo-EM image analysis of MS2nc@ApoF reassembly showed a mixed population of empty and cargo-filled MS2nc particles (**Figure 1B**). From 5,000 micrographs, we selected ∼240,000 particles and used icosahedral symmetry to produce a map of MS2nc at a resolution of 1.86 Å. Computational “particle subtraction” from cargo-containing 2D images revealed the ApoF inside, and standard single-particle analysis of the subtracted images yielded a 2.16 Å resolution structure of ApoF (**Figures 1C, S4**).

This level of resolution for ApoF is lower than the sub-2 Å resolution routinely achieved for the protein alone. This difference may stem from imperfect particle subtraction or a higher background due to the need for effectively thicker ice compared to what could be achieved with isolated proteins that are smaller than MS2nc. More sophisticated methods for accurate particle subtraction are being developed^38^ that may further improve resolution, though current approaches already deliver structures suitable for detailed molecular analysis. Importantly, the structure of nanocrate-packaged ApoF is indistinguishable from structures determined by conventional methods,^39,40^ demonstrating that the nanocrate environment does not induce conformational changes.

Having established that nanocrates can successfully encapsulate ApoF and that our proposed processing pipeline yields high-resolution reconstructions, we next tested the applicability of nanocrates with bovine thyroglobulin (Tg), a protein with well-documented preferred orientation problems in conventional cryo-EM. Previous high-resolution structures of Tg required detergents or other specialized procedures.^41-46^

Packaging Tg into MS2 nanocrates presented a challenge: the protein’s dimensions exceed the available inner diameter of the MS2 icosahedral shell. Rather than preventing encapsulation, this resulted in one lobe of Tg protruding through the MS2 shell (**Figure 1D**). This partial encapsulation still provided adequate protection from air-water interface effects, yielding a 2.94 Å resolution reconstruction with no denatured Tg particles observed in 2D classification.

The presence of both packaged and free thyroglobulin in the nanocrate preparation allowed for a direct comparison, revealing substantial advantages provided by encapsulation. Analysis of orientation distributions revealed that Tg from nanocrates achieved better angular sampling than non-encapsulated Tg from the same dataset (**Figure 2A, 2B**). More importantly, the nanocrate-derived structure achieved higher resolution (3.4 Å with C1 symmetry) compared to the structure (3.9 Å with C1 symmetry) obtained from equivalent numbers of free protein images, demonstrating that improved orientation distribution directly translates to better structural information.

**Figure 2.**
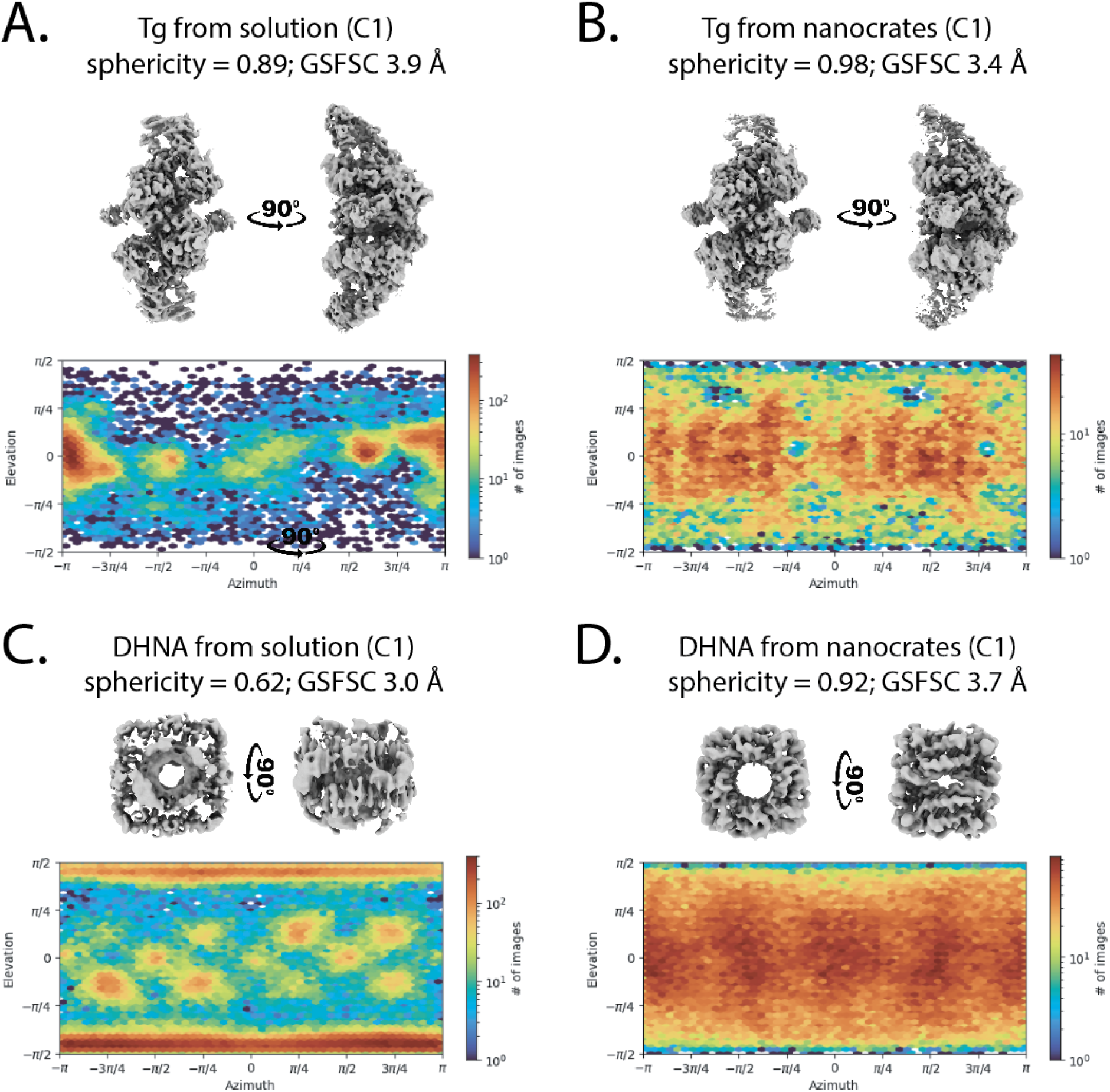
Nanocrate encapsulation eliminates preferred orientation. **A**. Cryo-EM map and orientation distribution plot of 32,904 Tg particles refined with C1 symmetry. **B**. Nanocrate-derived cryo-EM map and orientation distribution plot of 32,904 Tg particles refined with C1 symmetry. Note the improved resolution of the nanocrate-derived structure versus the solution map. Both reconstructions from **A** and **B** were derived from the same dataset, which contained both non-packed and MS2-packaged Tg molecules. **C**. Cryo-EM map and orientation distribution plot of 112,060 DHNA particles refined with C1 symmetry from the conventionally prepared cryo-EM grids. Despite relatively high GSFSC resolution, the map is unsuitable for model building due to low sphericity. **D**. Nanocrate-derived cryo-EM map and orientation distribution plot of 112,060 DHNA particles refined with C1 symmetry.

The Tg reconstruction revealed that the protein remained highly dynamic even within nanocrates, as evidenced by weaker density at distal regions (**Figure S5C**). Masked local refinement enabled better visualization of these dynamic regions (**Figure S5D-F**); however, full model building would require additional 3D classification and local filtering approaches.^41-46^ Thus, the use of nanocrates eliminated the preferred orientation artifacts that have historically complicated Tg structure determination, converting a problematic target into a routinely processable sample.

Finally, we applied MS2nc to the challenging cryo-EM target of 7,8-dihydroneopterin aldolase (DHNA), a barrel-shaped octameric protein assembly that adopts extreme preferred orientation on conventional grids. Our attempts to determine the DHNA structure by standard methods yielded highly anisotropic maps unsuitable for model building, despite achieving a GSFSC resolution of 2.29 Å (**Figure S7**). The orientation distribution plots revealed the severity of the problem: DHNA particles adopted essentially identical orientations, providing insufficient angular sampling for meaningful 3D reconstruction (**Figures 2C and 2D**). Since DHNA has an assembled molecular weight of 107 kDa and an expected diameter of 70 Å, we anticipated it would be readily packaged in MS2nc.

We observed MS2nc@DHNA particles containing multiple copies of DHNA in an ordered C5 symmetric arrangement (**Figure S6C**), allowing the use of a symmetry expansion workflow to obtain the final 3D reconstruction (**Figures S6F**). MS2nc@DHNA yielded a 2.80 Å resolution reconstruction (**Figure 1E**) with well-distributed particle orientations (**Figure 2D**) making it suitable for atomic model building.

From a methodological perspective, we found that one could not simply use the published structure of MS2 for the nanocrate density subtraction step. Instead, it was necessary to employ the experimental data for MS2nc obtained for each sample to calculate the shell density for subtraction. The nanocrate thereby provides a built-in resolution standard, determining the principal resolution limit of the dataset. It can also be used to tune dataset processing parameters such as coma, magnification anisotropy, per-particle CTF, Cs value, and pixel size.

In summary, nanocrates offer significant practical advantages for the high-resolution reconstruction of proteins by single-particle cryo-EM. The method improved the angular orientation of particles that suffered from preferred orientations by traditional plunge freezing methods. Using the same protocol, we observed significantly different encapsulation rates for the various cargos: 25% for MS2nc@ApoF, 30% for MS2nc@Tg, and 65% for MS2nc@DHNA. It proved unnecessary to modify the cargo loading protocol for each sample, as these variations in the extent of cargo loading did not hinder resolution as measured by gold-standard Fourier-shell correlation. Furthermore, unlike conventional methods requiring extensive optimization of grid preparation parameters, nanocrates enable standardized cryo-EM preparation since their consistent external surface properties, regardless of cargo, govern particle behavior during plunge freezing. This removes the need to re-optimize sample concentration, grid type, and blotting methods, providing significant time and cost savings while reducing consumption of expensive samples, consumables, and microscope time. The use of nanocrates also eliminates air-water interface effects through complete molecular shielding. While we expect that protein-cage interactions may occasionally cause problems, it should be possible to optimize the properties of the interior particle surface for specific specimens or to employ other self-assembling nanoparticle containers as nanocrates.

## Methods

### MS2 capsids expression and purification

MS2 capsids can be purified, disassembled, and reassembled by various methods,^29,30,47-49^ and the optimal protocol will depend on the equipment and standard procedures of each laboratory. In supplemental methods, we provide two such protocols – one from the Georgia Tech laboratory and another from NYSBC. Both provided approximately the same yield and quality of MS2 capsids: typically, 10-50 mg of purified MS2 VLPs were isolated from 1 L cell culture.

### Disassembly of MS2 VLPs

For simultaneous capsid disassembly and encapsulated RNA removal, 1 mL of glacial acetic acid was added dropwise over ∼30 seconds (without stirring) into a 0.5 mL solution of MS2 capsids (10 mg/mL) in 1x PBS. The solution was capped, mixed by inversion 2-3 times, and then incubated on ice for 20 minutes. The cloudy solution was then centrifuged at 6,600*g* for 10 minutes at 4^°^C to pellet out precipitated RNAs and any stochastically precipitated MS2 coat proteins. The clear supernatant, containing soluble disassembled MS2 coat protein dimers, was then removed by aspiration and loaded into a NAP-25 column (Cytiva) that had been pre-equilibrated with aqueous 1 mM acetic acid. MS2 dimers were eluted in 10 × 0.5 mL fractions of 1 mM acetic acid, which were immediately placed on ice. The protein concentration of each elution fraction was determined using the absorbance value at 280 nm on a NanoDrop instrument (using the theoretical calculation of 1.235 absorbance units for 1 mg/mL concentration, as determined from the MS2 primary sequence using the ExPASy ProtParam online tool, https://web.expasy.org/protparam/).^50^ The 3-4 most concentrated fractions were pooled and either used immediately for cargo packaging reactions or stored at -80^°^C for future use.

### Cargo packaging within reassembling MS2 VLPs

Packaging of cargo proteins within reassembling MS2 VLPs was achieved by first mixing the cargo protein of interest, water, and a 1/10^th^ volume of 10x TMK buffer (0.1 M Tris-HCl [pH 8.5], 80 mM KCl, 10 mM MgCl_2_)^47,51^ relative to the intended final reassembly solution volume in a 1.5 mL Eppendorf tube. Disassembled MS2 coat protein dimers in 1 mM acetic acid were subsequently added to complete the reassembly mixture. The Eppendorf tube was then capped and gently agitated at room temperature (∼25^°^C) with an end-over-end mixer for 3 hours. Following capsid reassembly, the sample mixture was stored at 4^°^C prior to shipping to NYSBC for cryo-EM imaging.

Several reassembly mixtures were prepared for each cargo molecule in which the final concentration of cargo protein monomers was varied in relation to a fixed concentration of MS2 coat protein monomers. In all cases, an abundance of MS2 coat proteins was included relative to cargo protein since each reassembling nanocrate requires 180 monomers to form a complete VLP shell. Consequently, the MS2 coat protein monomer to cargo monomer ratios (i.e., MS2:cargo ratios) were tested over a range of 12.5:1 up to 200:1 in nanocrate reassembly solutions. The specific MS2:cargo ratios used to collect the cryo-EM data presented in **Figures 1 and 2** are included in **Table S1**. An example reassembly mixture is presented in **Table S2**.

### Negative staining and EM data acquisition

EM grids (Ted Pella Inc, Carbon Type-B copper 300 mesh) were plasma cleaned using a H_2_/O_2_ gas mixture for 30 s in a Solarus Plasma Cleaner 950 (Gatan). The VLP sample (3 µL, 0.1 mg/mL in PBS) was applied to the grid and allowed to adsorb for 30 s before blotting away excess liquid. This was followed by two cycles of washing with deionized (Milli-Q) water, blotting, and staining with 2% (w/v) uranyl acetate for 45 s before final blotting. Negatively stained grids were imaged using a Hitachi-7800 transmission electron microscope at an accelerating voltage of 100 keV, a nominal magnification of 120,000x (corresponding to a pixel size of 1.8 Å) and defocus ranging from -2.0 to -3.0 µm.

### Cryo-EM sample preparation and data acquisition

Unless otherwise specified, the MS2nc@cargo samples were concentrated 10-fold using spin concentrators with a 100 kDa molecular weight cut-off. The sample was applied to plasma-cleaned UltrAuFoil 1.2/1.3 grids (H_2_/O_2_ gas mixture for 7 s in a Solarus Plasma Cleaner 950 (Gatan)) and imaged on a TFS Titan Krios instrument G2 equipped with a Gatan K3 camera and a Gatan Bioquantum energy filter. Data were acquired using Leginon^52,53^ in counting mode with a calibrated pixel size of 0.832 Å and a 20 eV energy slit. The total dose was 55 e^-^/A^2^ per movie, split into 50 frames. Movie stacks were motion corrected and dose-weighted with motioncorr2^54^, and the resulting images were imported into CryoSPARC^55^ for processing.

### Data processing

Data processing was performed using standard SPA tools in two major steps. First, the dataset was processed to achieve high-resolution reconstruction of the nanocrate, followed by particle subtraction. Second, the subtracted stack was processed to reconstruct the cargo proteins. The overall processing workflow is summarized in **Figure S2**, and detailed descriptions of individual cargoes are provided in Supplemental Methods. Plots in 1C, 1D, 1E, and sphericity values were generated by processing final refinements using the 3DFSC server.^6^

## Acknowledgements

We thank Bridget Carragher for inspiration and support, Christopher JF Cameron and Joshua Mendez for invaluable discussions, Aaron Owji for advice with data processing, and Christina Bourne for providing the DHNA sample. This work was supported by the Simons Foundation (SF349247) for structural studies at the Simons Electron Microscopy Center at the New York Structural Biology Center, and by the National Institutes of Health (R01 AI148382, R01 CA247484).

## Supplemental methods

### MS2 expression and purification – method #1 (Georgia Tech)

The pCDF:MS2 plasmid encoding the MS2 coat protein gene was transformed into chemically competent BL21(DE3) *E. coli* cells (New England Biolabs), and positive transformants were selected by plating cells onto 2YT agar plates containing a final concentration of 50 µg/mL streptomycin. After overnight incubation at 37^°^C, transformant cells were gently scraped from a single colony with a sterile pipette tip and were inoculated into 500 mL of 2YT or SOB medium supplemented with 50 µg/mL streptomycin in a 2 L baffled Erlenmeyer flask. The culture flask was then grown to saturation by incubation at 37^°^C for 20-24 hours in a shaking incubator set to a rotational speed of 180 rpm. Cells were subsequently harvested from the culture volume by centrifugation at 8,000 rpm for 15 minutes at 4^°^C in a JLA-16.250 rotor (Beckman Coulter), and the resulting cell pellets were stored at -80^°^C until purification.

Purification of MS2 VLPs was performed by first resuspending cell pellets from an entire 0.5 L expression batch in 50 mL of 50 mM potassium phosphate buffer (pH 7.0). The resuspended cell slurry was transferred to a 100 mL glass beaker. The cells were lysed via sonication in an ice/water bath using a QSonica Q500 sonicator equipped with a 0.5” probe tip (5-second sonication pulses with 5-second rest between pulses at an instrument amplitude value between 45-55%, 120 pulses in total). The cell lysate was then clarified by centrifugation at 14,000 rpm for 15 minutes at 4^°^C in a JA-17 rotor (Beckman Coulter). VLPs were subsequently precipitated from the clarified lysate by adding solid ammonium sulfate to a final concentration of 0.265 g/mL, dissolving the ammonium sulfate solids at room temperature, and then incubating the mixture on an end-over-end mixer at 4^°^C for 1 hour. Precipitated solids were collected by centrifugation at 14,000 rpm for 15 minutes at 4^°^C in a JA-17 rotor and resuspended in 8 mL of fresh 50 mM potassium phosphate buffer (pH 7.0). Residual hydrophobic contaminants were removed by measuring the volume of the VLP solution (which usually expanded to 10-12 mL following solubilization of precipitated solids) and then adding an equal volume of an organic solution consisting of *n*-butanol and chloroform in a 1:1 ratio to the aqueous VLP solution. The aqueous/organic mixture was vigorously agitated for 1 minute, then the respective aqueous and organic layers were resolved by centrifugation at 14,000 rpm for 10 minutes at 4^°^C in a JA-17 rotor. The upper aqueous layer was gently removed by aspiration and then layered on top of 10-40% sucrose gradients (∼3 mL VLP solution per 40 mL gradient tube) prepared in 50 mM phosphate buffer (pH 7.0). Gradient sedimentation was performed by ultracentrifugation at 28,000 rpm for 4 hours at 4^°^C in a SW32 rotor (Beckman Coulter).

VLPs were visualized by bottom-up white light illumination of sucrose gradient tubes and were carefully removed by aspiration with a syringe. Isolated VLPs were then concentrated by ultracentrifugation at 68,000 rpm for 2 hours at 4^°^C in a Type 70 Ti rotor (Beckman Coulter), and the resulting supernatant was decanted. The pelleted VLPs in each ultracentrifuge tube were gently resuspended in 3-5 mL of fresh 1x PBS buffer, sterile filtered through a 0.2 µm PES syringe filter, and stored at 4^°^C (short-term) or -80^°^C (long-term). Before storage, the concentration of purified MS2 VLPs was determined using a Bradford assay (Pierce) with BSA serving as the protein standard.

### MS2 expression and purification – method #2 (NYSBC)

The pCDF:MS2 plasmid was transformed into *E. coli* BL21(DE3) Gold cells (Agilent) and plated onto a Luria-Bertani (LB) agar plate containing 50 μg/mL streptomycin. A single colony was used to inoculate 20 mL of LB medium supplemented with the same antibiotic, and the culture was incubated overnight at 37^°^C with shaking at 225 rpm. This preculture was then used to inoculate 2 L of LB medium containing 50 μg/mL streptomycin and 0.01% Antifoam 204 and was grown at 37^°^C in a LEX bioreactor (Epiphyte Three Inc.) until an optical density of 0.8 measured at 600 nm was reached. Protein expression was induced by the addition of 1 mM isopropyl β-D-thiogalactopyranoside (IPTG) with overnight shaking at 25^°^C. Cells were harvested by centrifugation at 4,000*g* for 20 minutes and stored at −80^°^C until needed.

A cell pellet from 2 L of culture (approximately 9.5 g) was resuspended in 60 mL of 0.1 M potassium phosphate buffer pH 7.5 and lysed by sonication for 10 min (65% amplitude, 5 seconds on / 5 seconds off). Cell debris was removed by centrifugation at 27,000*g* for 15 minutes at 4^°^C. Ammonium sulfate was added to the supernatant to a final concentration of 0.265 g/mL and stirred until fully dissolved. The mixture was rotated for 1 hour at 4^°^C and precipitated proteins were collected by centrifugation at 27,000*g* for 15 minutes at 4^°^C. The resulting pellet was gently resuspended in 16 mL of 0.1 M potassium phosphate buffer pH 7.5 and incubated overnight at 4^°^C with gentle rotation. An equal volume of a 1:1 n-butanol/chloroform mixture was added, and the solution was vigorously mixed for 1 minute. After centrifugation at 27,000*g* for 10 minutes at 4^°^C, the upper aqueous phase was collected, diluted 50-fold with 10 mM Tris-HCl buffer pH 8.0, and filtered through a 0.4 μm filter. The clarified sample was loaded onto a Mono Q 10/100 GL anion exchange column (Cytiva) and MS2 protein was eluted using a NaCl gradient in 10 mM Tris(hydroxymethyl)aminomethane (Tris) pH 8.0. Fractions eluting around 500 mM NaCl were pooled, concentrated to 5 mL, and further purified using a HiLoad 16/60 Superdex 200 prep-grade column (Cytiva) equilibrated with 1x PBS pH 7.4. Purified protein fractions were pooled, concentrated to 10 mg/mL (measured by Bradford assay), and stored at −80^°^C.

### MS2 capsid and MS2nc empty nanocrate cryo-EM data processing

Processing for empty nanocrates is summarized in **Figure S3**. Motion correction was performed with MotionCor2 to generate motion-corrected and dose-weighted images; all subsequent zrocessing was done using cryoSPARC v4.6.2. CTF estimation was done using the Patch CTF module with default settings. Micrographs were curated based on a CTF resolution cutoff of 7 Å and a defocus range of 0.7–2.0 µm, resulting in 4,357 accepted and 443 rejected micrographs. Particles were picked using a single high-quality template view of MS2 with a particle diameter of 400 Å and 100 expected particles per micrograph. A total of 259,778 particles were extracted using a 480-pixel box size at a pixel size of 0.826 Å. The initial cleanup involved 2D classification with default settings, followed by the selection of sharp icosahedral classes (234,080 particles). *Ab initio* reconstruction and non-uniform refinement with enforced icosahedral symmetry, local CTF refinement, and EWS correction yielded a GSFSC 1.74 Å resolution map. The final resolution MS2nc dataset is the highest resolution of any MS2 structure reported to date, and the resulting density had resolved waters, so we proceeded to build an atomic model.

Coordinates from PDBID 2IZM with all non-protein atoms removed were used as an initial model. The asymmetric unit was rigid-body fitted into the map using UCSF Chimera. Iterative rounds of real space minimization and ADP refinement in Phenix^56^ and manual inspection and adjustment in Coot^57^ were performed. Only minor shifts in the rotamers of a few side chains were made from the initial model. Map peaks for water molecules were visible (**Fig S3E**). Waters were added to the model using the “Find Waters” feature in Coot for peaks above 2.0 σ. Added waters were curated by visual inspection, with additional waters added or deleted manually based on observed map peaks and the presence of neighboring hydrogen bonding partners. The final model includes an asymmetric unit of three chains, residues 1-129. Residues 1 and 13-15 of each chain are less resolved, but backbone density is visible. All other residues are well ordered with clear sidechain density (**Figure S3E**). The model includes 228 water molecules.

During this work, we generated a reconstruction of the MS2 capsid prior to the disassembly/reassembly process. The resulting reconstruction had equivalent map quality to the reassembled empty nanocrate. For completeness, we also built an atomic model into this map. The coordinates from the nanocrate model refinement, including the built waters, were used as a starting model. An initial round of rigid body, global real-space minimization, and ADP refinement was performed in Phenix. Minimal rotamer adjustments were made manually in Coot based on visual inspection, followed by iterative rounds of real space minimization and ADP refinement in Phenix and manual inspection and adjustment in Coot to improve the geometry and verify the placement of water molecules. Based on manual inspection, an additional ∼40 waters were added and ∼5 waters deleted relative to the nanocrate starting model. We did not analyze these differences in water structure further, as it is outside the scope of this work.

Cryo-EM maps and models were deposited to PDB/EMDB as follows:

|  |  |  |
| --- | --- | --- |
| 9Q1B<br>EMD-72122 | / | MS2 bacteriophage coat protein after reassembly as a nanocrate (MS2nc) with no cargo and with waters modeled |
| 9Q1D<br>EMD-72124 | / | MS2 bacteriophage coat protein with waters modeled |
| EMD-72123 |  | Mouse apoferritin encapsulated in an MS2-nanocrate: MS2nc@ApoF |
| EMD-72176 |  | Porcine thyroglobulin encapsulated in an MS2-nanocrate: MS2nc@Tg |
| EMD-72177 |  | Dihydroneopterin aldolase encapsulated in an MS2-nanocrate: MS2nc@DHNA |

### MS2nc@ApoF cryo-EM data processing – particle subtraction and unsubtracted refinement

Processing of ApoF from nanocrates is summarized in **Figure S2**, and results are shown in **Figure S4**. Motion correction was performed using MotionCor2 to generate motion-corrected and dose-weighted images; all subsequent processing was carried out using cryoSPARC v4.6.2. CTF estimation was done using the Patch CTF module with default settings. Micrographs were curated based on a CTF resolution cutoff of 4.5 Å and a defocus range of 0.7–2.0 µm, resulting in 2,860 accepted and 2,036 rejected micrographs. Particles were picked using a single high-quality template view of MS2 with a particle diameter of 400 Å and 100 expected particles per micrograph. A total of 348,343 particles were extracted using a 480-pixel box size at a pixel size of 0.826 Å. Initial cleanup involved 2D classification with default settings, followed by selection of sharp icosahedral classes (239,460 particles). *Ab-initio* reconstruction and homogeneous refinement with enforced icosahedral symmetry, local CTF refinement and EWS correction produced a GSFSC 1.86 Å map. Particle subtraction was performed using particles from the final refinement. The resulting cage-subtracted stack was subjected to further 2D classification, and 69,412 particles containing ApoF were selected. ApoF-containing classes (∼30% of the initial stack) were chosen for initial model generation and homogeneous refinement with octahedral symmetry enforced, resulting in a GSFSC 2.16 Å resolution map of ApoF (**Figure S4C**). Displayed at a lower threshold, we can see a noise shell around the ApoF, corresponding to the radially averaged out partially assembled MS2nc cage around ApoF molecules that can be seen in some 2D classes. Using the alignment information from the particle-subtracted reconstruction, we reconstructed particles from the unsubtracted stack without any refinement. The resulting map (**Figure S4E**) has achieved a GSFSC resolution of 2.19 Å. When displayed at the lower threshold, a second noise shell can be seen around the ApoF density, corresponding to the MS2nc density that is radially averaged out.

### MS2nc@Tg cryo-EM data processing – particle subtraction, local refinement, 3D classification and non-nanocrate processing

Processing of Tg from nanocrates is summarized in **Figure S5**. Motion correction was performed with MotionCor2 to generate motion-corrected and dose-weighted images; all subsequent processing was done using cryoSPARC v4.6.2. CTF estimation was done using the Patch CTF module with default settings. Micrographs were curated based on a CTF resolution cutoff of 4.0 Å and a defocus range of 0.7–2.0 µm, resulting in 7,549 accepted and 1,645 rejected micrographs. Particles were picked using a single high-quality template view of MS2nc with a particle diameter of 400 Å and 100 expected particles per micrograph. A total of 681,983 particles were extracted using a 480-pixel box size at a pixel size of 0.826 Å. Initial cleanup involved 2D classification with default settings, followed by selection of sharp icosahedral classes (541,146 particles). *Ab-initio* reconstruction and homogeneous refinement with enforced icosahedral symmetry yielded a GSFSC 2.02 Å map. After local CTF refinement and EWS correction, further homogeneous refinement improved the resolution to 1.92 Å. Particle subtraction was performed using particles from the final refinement. The resulting cage-subtracted stack was subjected to further 2D classification, and 131,048 particles containing Tg were selected. Homogeneous refinement with default settings and C2 symmetry resulted in a GSFSC 2.92 Å resolution map (**Figure S5C**). Symmetry expansion followed by masked local refinement of the asymmetric unit resulted in a GSFSC 2.80 Å resolution map, with a better-resolved distal part of the Tg map within the masked area (**Figure S5E**). At the same time, the distal part of the Tg map that was outside the mask became significantly distorted, suggesting that Tg molecules within the nanocrystals remained highly dynamic.

From the same dataset, we processed Tg particles that were not incorporated into nanocrates. These were picked using the template picker, 2D classified to retain only Tg-containing classes, further classified in 3D using *ab-initio* with 3 classes, and resulted in a final stack of 32,904 particles. Homogeneous refinement with C1 symmetry produced a GSFSC 3.87 Å resolution map, which was used to compare to the MS2nc@Tg derived map (**Figure 2A, 2B**).

### MS2nc@DHNA cryo-EM processing –particle subtraction and custom symmetry expansion

Processing of DHNA from nanocrates is summarized in **Figure S6**. Motion correction was performed with MotionCor2 to generate motion-corrected and dose-weighted images, all subsequent processing was done using cryoSPARC v4.6.2. CTF estimation was done using the Patch CTF module with default settings. Micrographs were curated based on a CTF resolution cutoff of 4.5 Å and a defocus range of 0.7–2.0 µm, resulting in 6,360 accepted and 4,160 rejected micrographs. Particles were picked using a single high-quality template view of MS2 with a particle diameter of 400 Å and 100 expected particles per micrograph. A total of 252,070 particles were extracted using a 480-pixel box size at a pixel size of 0.826 Å. Initial cleanup involved 2D classification with default settings, followed by selection of sharp icosahedral classes (149,058 particles). *Ab-initio* reconstruction and homogeneous refinement with enforced icosahedral symmetry yielded a 2.34 Å map. After local CTF refinement and EWS correction, further homogeneous refinement improved the resolution to 2.24 Å. Particle subtraction was performed using particles from the final refinement. The resulting cage-subtracted stack was subjected to further 2D classification, and 84,838 particles containing multiple DHNA copies were selected. A series of *ab-initio* classifications and homogeneous refinements led to a final subset of 25,543 particles displaying well-resolved DHNA packing (∼10 Å resolution). Non-uniform refinement with a new reference and C5 symmetry enforcement yielded a GSFSC cargo map at 6 Å resolution. An additional non-uniform refinement with symmetry relaxation (marginalization) was applied to produce the final map used for sub-volume extraction. Local refinement was done following the cryoSPARC case study on encapsulated ferritin processing (see https://guide.cryosparc.com/processing-data/tutorials-and-case-studies/case-study-end-to-end-processing-of-encapsulated-ferritin-empiar-10716), beginning with five DHNA regions selected from the cargo map. Custom masks were used for sub-volume extraction with volume alignment tools in Chimera, enabling downstream local refinements and 3D classifications. Sub-volumes were aligned to a D4 symmetry axis, and 127,715 particles (5x expanded from the final cargo stack) were subjected to 3D classification into 10 classes. Selected particles (112,060) underwent local refinement with D4 symmetry, yielding a final map at 2.80 Å resolution.

### Non-nanocrate DHNA cryo-EM data processing

Processing of DHNA from solution is summarized in **Figure S7**. Motion correction was performed with MotionCor2 to generate motion-corrected and dose-weighted images, all subsequent processing was done using cryoSPARC v4.6.2. CTF estimation was done using the Patch CTF module with default settings. Micrographs were curated based on a CTF resolution cutoff of 4.5 Å and a defocus range of 0.7–2.0 µm, resulting in 187 accepted and 211 rejected micrographs. Particles were manually selected to generate an initial template, and then the template picker was used with a particle diameter of 70 Å and an expected number of 4000 particles per micrograph. A total of 571,862 particles were extracted using a 256-pixel box size at a pixel size of 0.829 Å. Initial cleanup involved 2D classification with default settings, followed by selection of sharp, well-centered classes (218,565 particles). *Ab-initio* reconstruction and homogeneous refinement C1 symmetry produced a GSFSC 2.73 Å resolution highly anisotropic map that was used to compare with the nanocrate-derived DHNA map (**Figure 2C, 2D**). Homogeneous refinement with D4 symmetry also produced a highly anisotropic map that wasn’t suitable for model building despite the GSFSC 2.29 Å resolution.

## Supplemental Figures and Tables

**Table S1.** Protein compositions of reassembly mixtures for the MS2nc@ApoF, MS2nc@Tg, and MS2nc@DHNA samples presented in the main text.

| Cargo Protein | MS2 monomer:<br>cargo monomer<br>ratio* | MS2 cage: cargo<br>ratio** | MS2 monomer<br>final conc. (μM) | Cargo monomer<br>final conc. (μM) |
| --- | --- | --- | --- | --- |
| ApoF (24-mer) | 25:1 | 3.33:1 | 125 | 5 |
| Tg (2-mer) | 75:1 | 0.83:1 | 125 | 1.67 |
| DHNA (8-mer) | 12.5:1 | 0.56:1 | 125 | 10 |

\* Ratio of MS2 coat protein monomer to cargo protein monomer
\*\* Ratio of an assembled MS2 cage (180 CPs) to cargo in its quaternary state

**Table S2.** Example reassembly mixture.

| Solution Component | Stock<br>concentration | Final<br>concentration | Volume added<br>(μL) |
| --- | --- | --- | --- |
| MS2 solution (disassembled) | 162 μM | 125 μM | 463.0 |
| Apo ferritin monomer | 123 μM | 5 μM | 24.4 |
| TMK buffer | 10x | 1x | 60.0 |
| Water | – | – | 52.6 |
| *Total volume* |  |  | *600 μL* |

**Figure S1.**
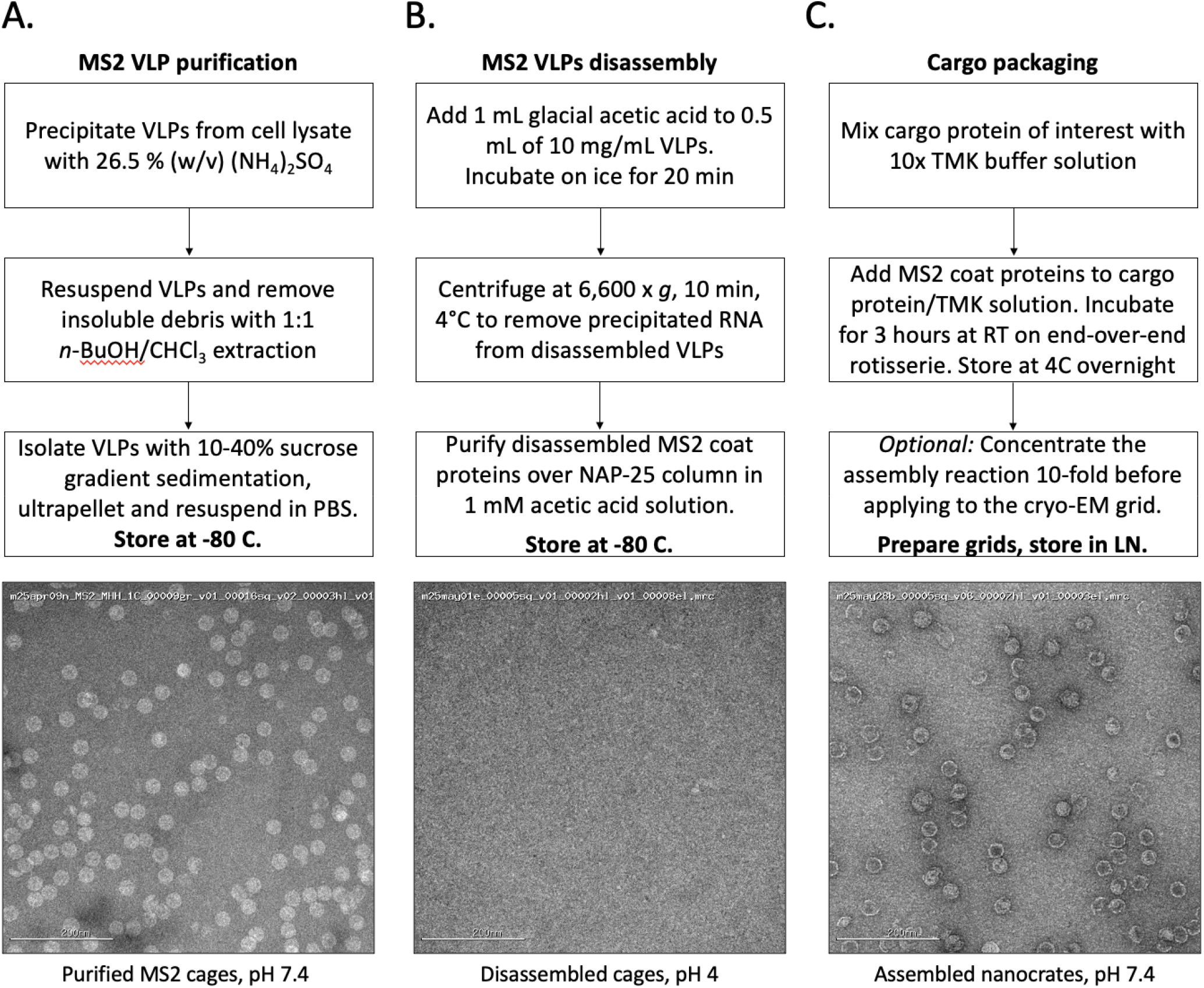
MS2 purification, disassembly and cargo packaging workflow.

**Figure S2.**
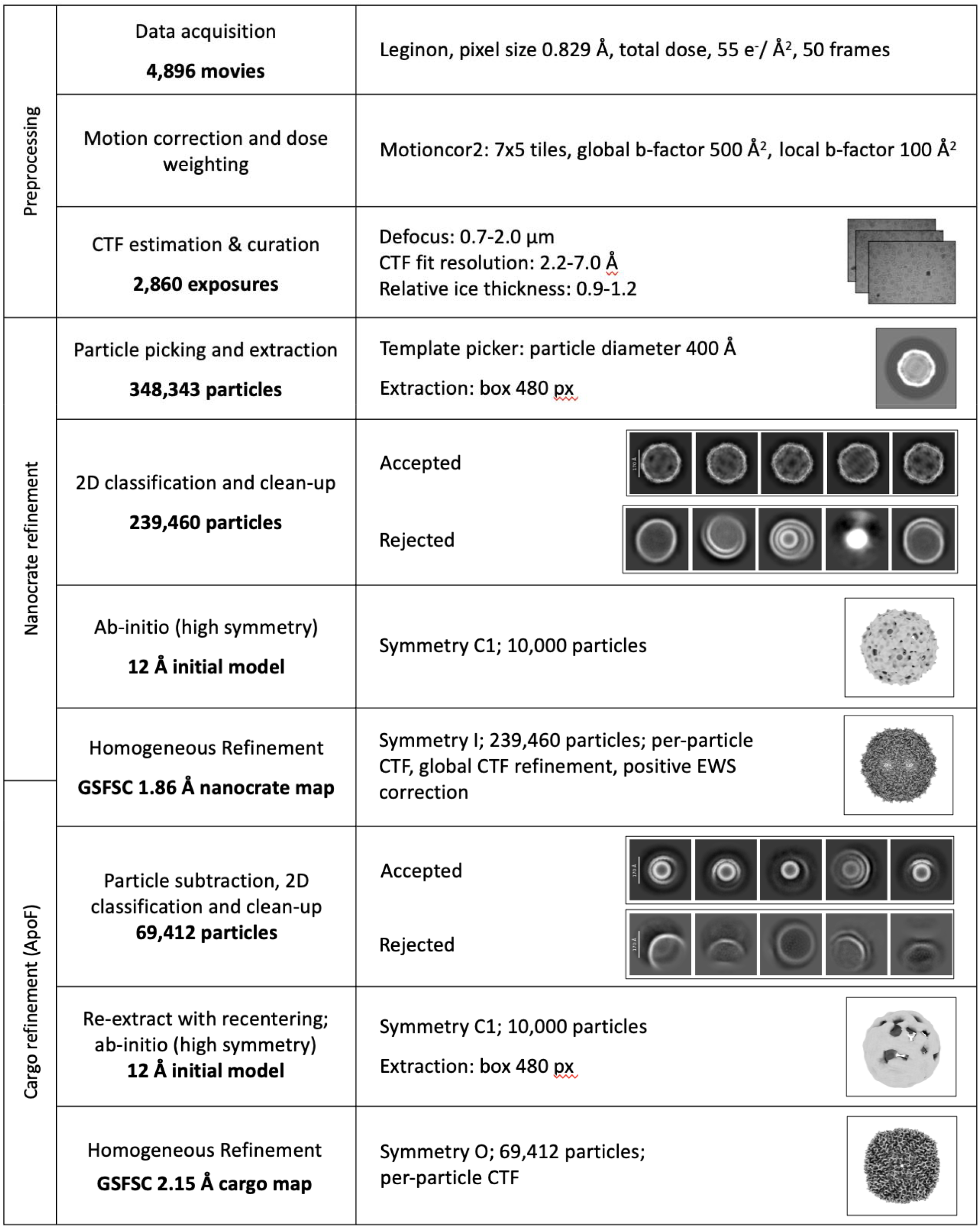
Caged particle reconstruction processing workflow.

**Figure S3.**
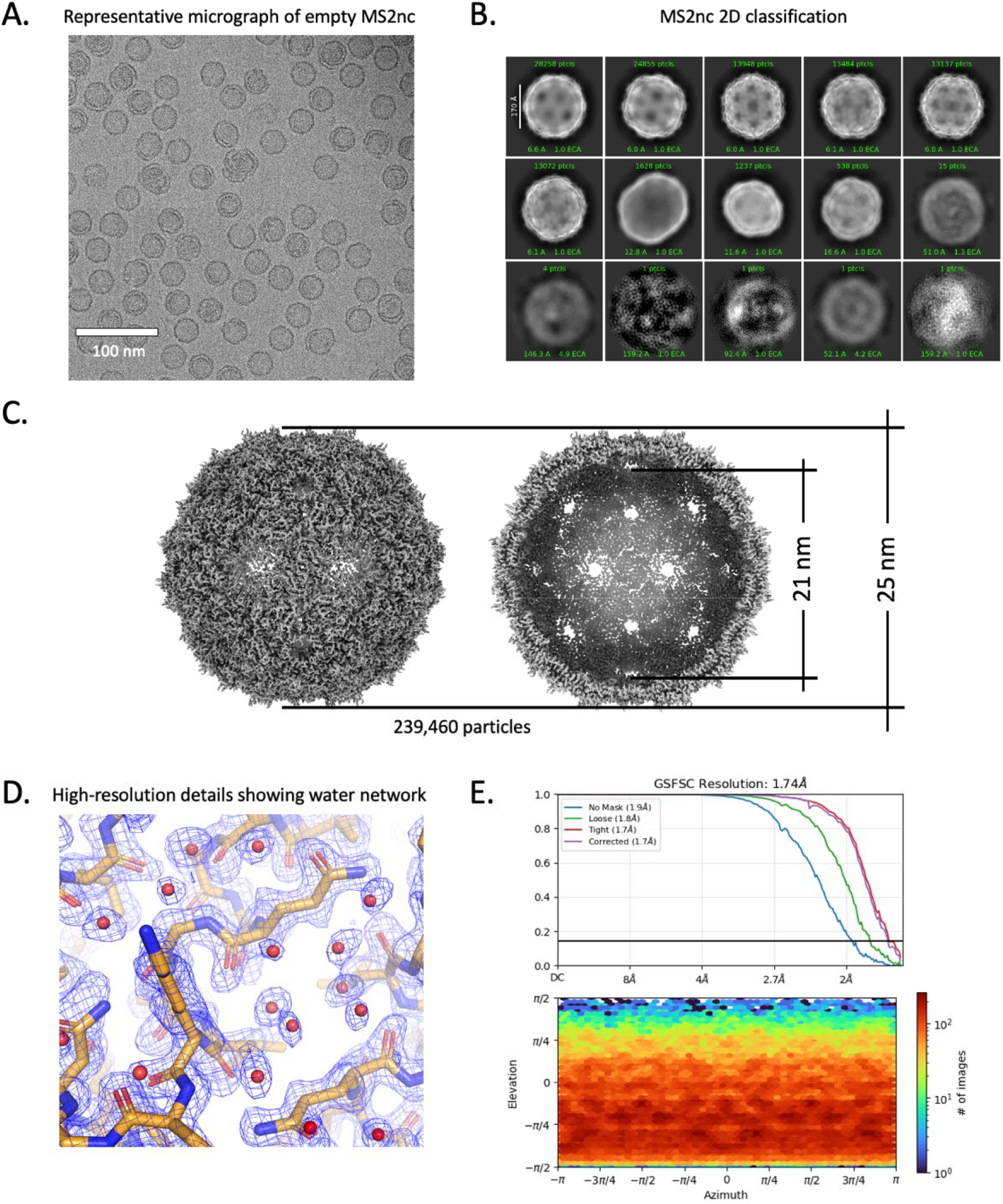
Empty MS2-nanocrate processing details. **A**. Representative micrograph of empty MS2-nanocrates. **B**. 2D classification of MS2nc without particle subtraction. **C**. refined map with I symmetry, MS2 nanocrate outer diameter = 25 nm, inner diameter = 21 nm, opening diameter of a 5-fold or 6-vertex pores = ∼1.5 nm. **D**. Detailed view of a selected area of the MS-nanocrate map (shown as blue mesh), showing high-resolution resolvability of sidechains and coordinated water. Modeled protein is displayed in stick representation with carbon in tan, oxygen in red, and nitrogen in blue. Waters are depicted as red spheres. **E**. GSFSC plot showing 1.74 Å resolution and viewing direction distribution plot, showing isotropic orientation distribution.

**Figure S4.**
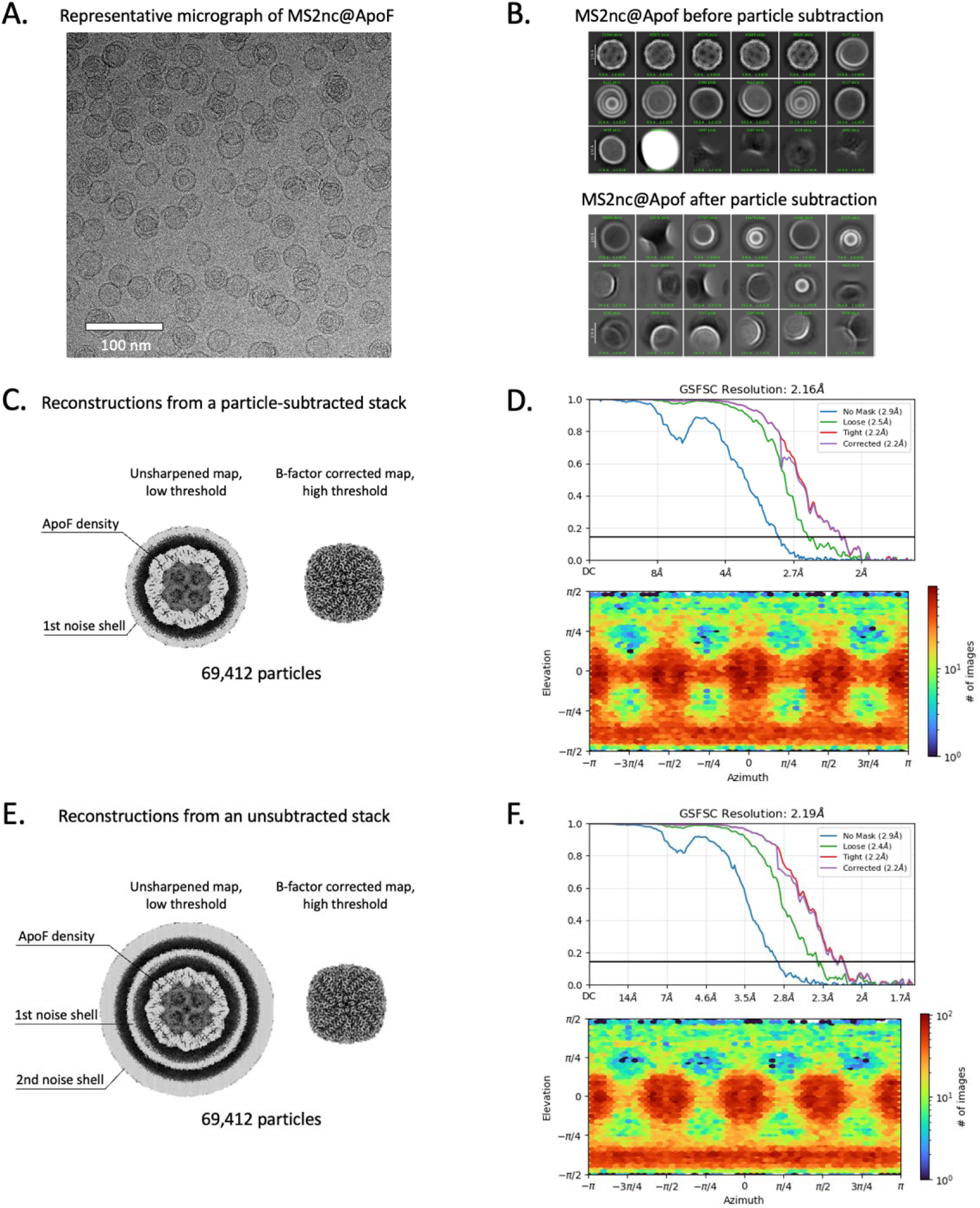
MS2nc@ApoF processing details. **A**. Representative micrograph. **B**. 2D classification before (top) and after nanocrate density subtraction. **C**. Refined map with O symmetry. **D**. GSFSC plot showing 2.16 Å resolution and viewing direction distribution plot, showing isotropic orientation distribution. **E**. Refinement with static mask, but without particle subtraction, using smaller box size and displayed at low threshold – nanocrate densities are averaged out into a uniform noise shell around the ApoF map. **F**. GSFSC plot showing 2.19 Å resolution and viewing direction distribution plot, showing isotropic orientation distribution.

**Figure S5.**
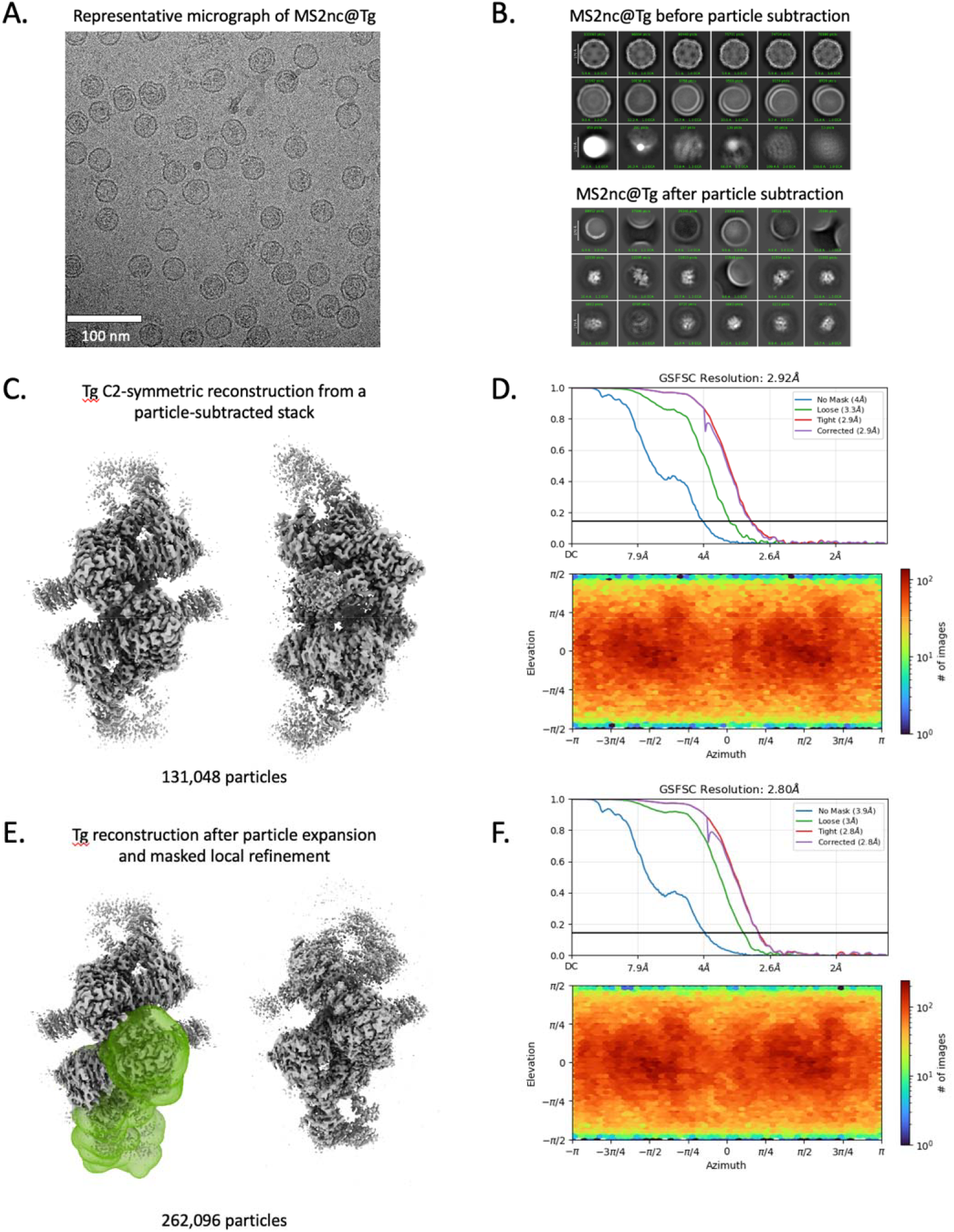
MS2nc@Tg processing details. **A**. Representative micrograph. **B**. 2D classification before (top) and after nanocrate density subtraction. **C**. Refined map with C2 symmetry. **D**. GSFSC plot showing 2.92 Å resolution and viewing direction distribution plot, showing isotropic orientation distribution. **E**. (*left*) Tg map with a mask region highlighted; (*right*) C1 reconstruction after local refinement of the masked region with corresponding GSFSC and viewing distribution plots **F**.

**Figure S6.**
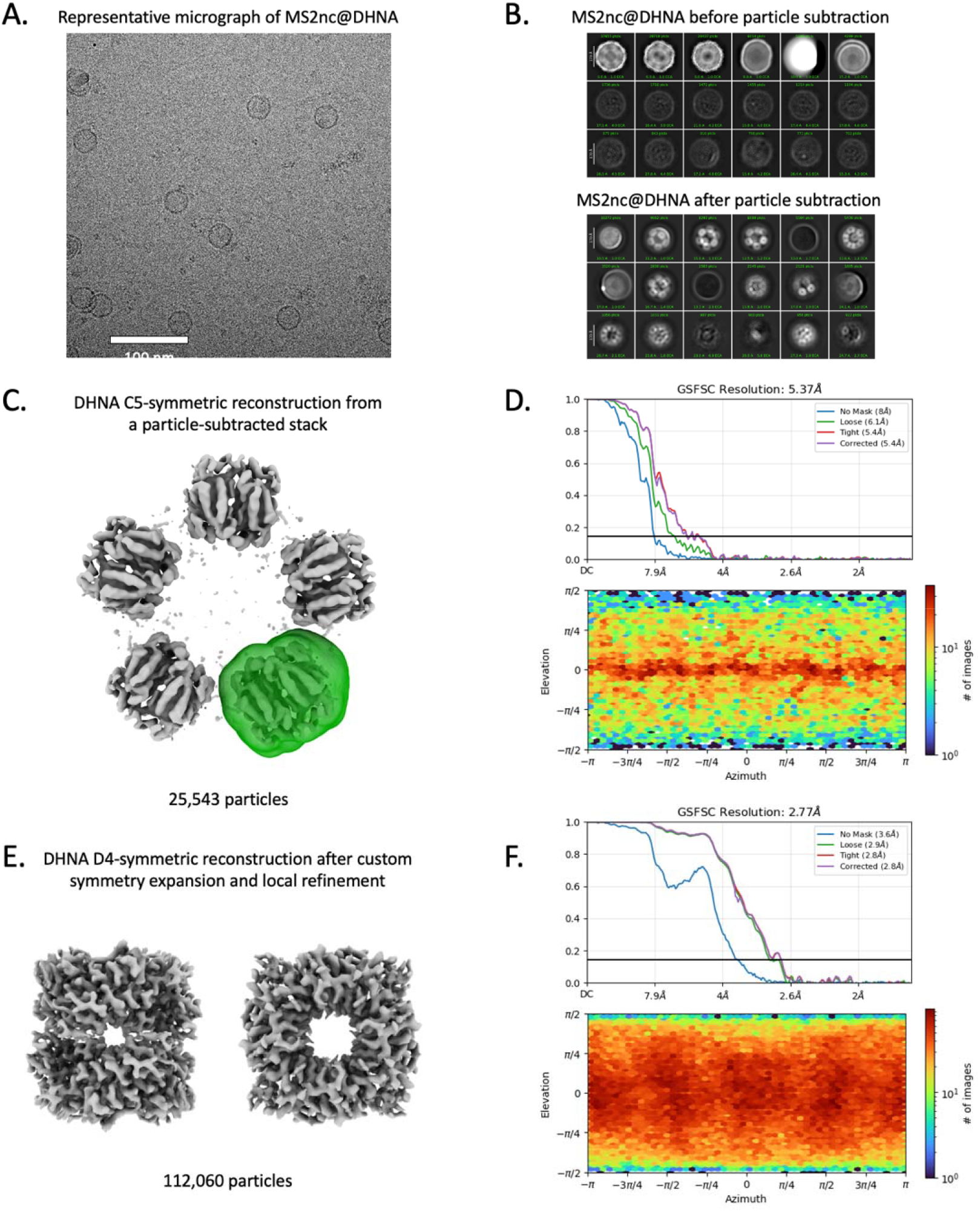
MS2nc@DHNA processing details. **A**. Representative micrograph. **C**. 2D classification before (*top*) and after nanocrate density subtraction. **C**. Refined map with C5 symmetry. A single DHNA copy was selected as a reference volume (masked in green) for symmetry expansion processing. **D**. GSFSC plot showing 5.375 Å resolution and viewing direction distribution plot, showing orientation distribution predominantly along the equator. **E**. Refined DHNA map after symmetry expansion and local refinement with D4 symmetry. **F**. GSFSC and viewing distribution plots corresponding to the reconstruction in **E**.

**Figure S7.**
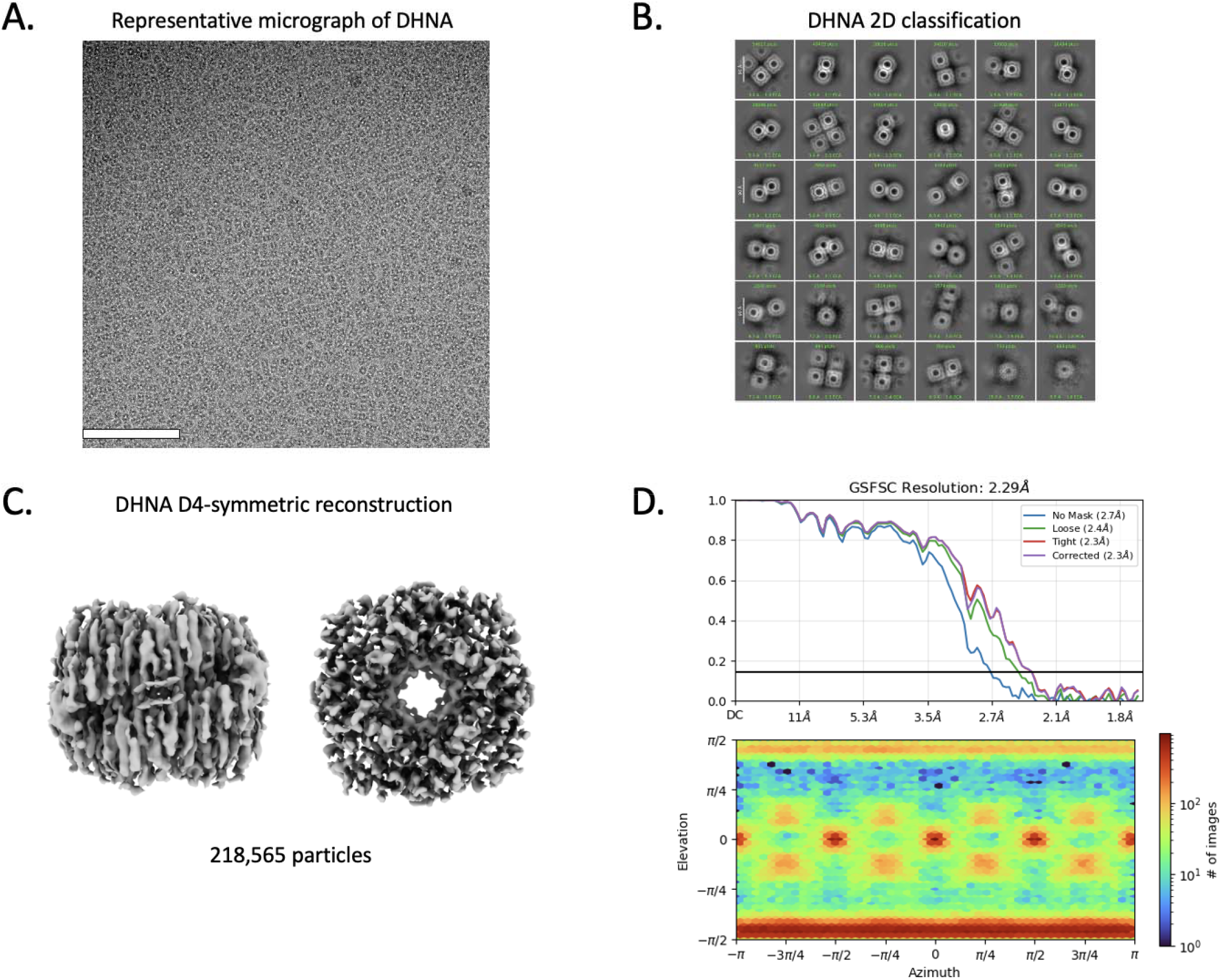
DHNA processing without nanocrates. **A**. Representative micrograph. **B**. 2D classification showing exclusively top views. **C**. Refined map with D4 symmetry enforced. The map is highly anisotropic and stretched in the direction of the preferred view, making it unsuitable for model building **D**. GSFSC plot showing 2.29 Å resolution and viewing direction distribution plot, showing highly preferred orientation.

**Cryo-EM data collection, refinement, and validation statistics for MS2 capsid and empty MS2nc**

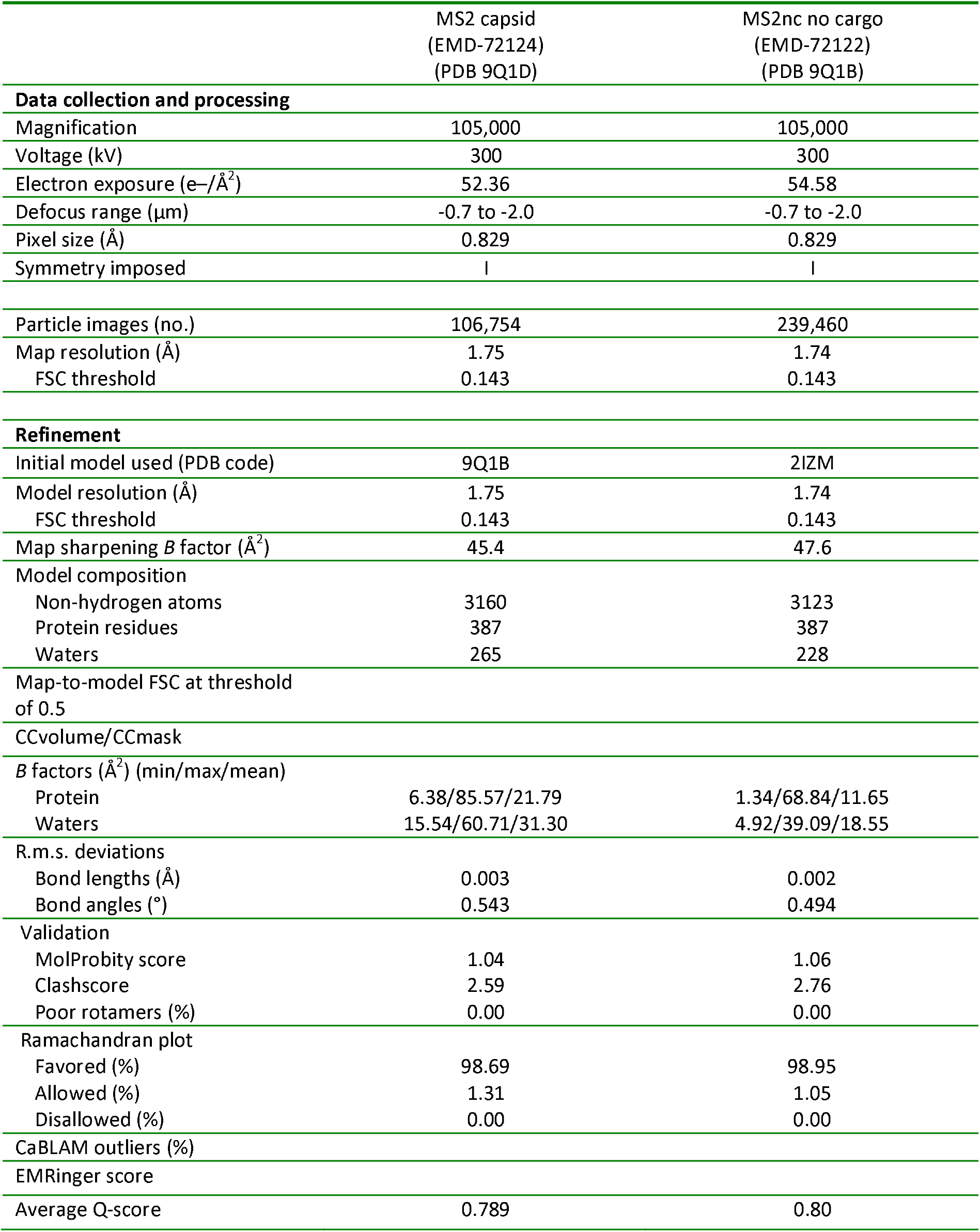

## Notes

### Competing Interest Statement

The authors have declared no competing interest.

